# CaV3.1 channel pore pseudo-symmetry revealed by selectivity filter mutations in their domains I/II

**DOI:** 10.1101/626069

**Authors:** E. Garza-López, A. Aldana, A. Darszon, T. Nishigaki, I. López-González

## Abstract

There is growing evidence indicating that the pore structure of voltage-gated ion channels (VGICs) influences gating besides their conductance. Regarding low voltage-activated (LVA) Ca^2+^ channels, it has been demonstrated that substitutions of the pore aspartate (D) by a glutamate (D-to-E substitution) in domains III and IV alter channel gating properties such as a positive shift in the channel activation voltage dependence. In the present report, we evaluated the effects of E-to-D substitution in domains I and II on the Ca_V_3.1 channel gating properties. Our results indicate that substitutions in these two domains differentially modify the gating properties of Ca_V_3.1 channels. The channel with a single mutation in domain I (DEDD) presented slower activation and faster inactivation kinetics and a slower recovery from inactivation, as compared with the WT channel. In contrast, the single mutant in domain II (EDDD) presented a small but significant negative shift of activation voltage dependence with faster activation and slower deactivation kinetics. Finally, the double mutant channel (DDDD) presented intermediate properties with respect to the two single mutants but with fastest deactivation kinetics. Overall, our results indicate that single amino acid modification of the selectivity filter of LVA Ca^2+^ channels in distinct domains differentially influence their gating properties, suggesting a pore pseudo-symmetry.

**Statement of significance:** Previous reports of low voltage-activated (LVA) Ca^2+^ channels have demonstrated that pore aspartate (D) in domains III and IV equally modulates the channel gating properties, supporting a hypothesis of pore symmetry in LVA Ca^2+^ channels. In the present report, we evaluated the effects of glutamate (E)-to-D pore substitution in domains I and II on the Ca_V_3.1 channel gating properties. Our results indicate that substitutions in these two domains differentially modify the gating properties of Ca_V_3.1 channels, therefore suggesting a pore pseudo-symmetry in them. Interestingly, our pore mutations affect inactivation of Ca_V_3.1 Ca^2+^ channels indicating a selectivity filter contribution to this process similar to the recent proposed paradigm for high voltage-activated Ca^2+^ channels.

## 1. Introduction

In general, voltage-gated ion channels (VGICs) are constituted at least by two basic domains: the voltage-sensing domain (VSD) formed by the first four transmembrane segments (S1-S4), where the S4 bears the positively charged residues; and the ion-conducting pore domain, which includes the S5-S6 plus the P-loop [1]. Previous papers have supported the idea that these domains independently regulate two different ion channel processes, the voltage-dependent channel gating and the ionic conductance (selectivity and permeability), respectively. In fact, the current model based on the K^+^ channel activation proposes that the membrane depolarization-induced outward movement of the VSD S4 segment leads to the channel gate opening by pulling the S4-S5 linker and bending the S6 segments [2]. The classical view of VGICs implies that the amino acid residues of one domain do not have a strong influence on the other domain’s function. However, there are several published papers suggesting a different perspective. Mutations within pore residues of potassium [3,4], sodium [5,6], cyclic nucleotide-gated (CNG) [7] or high voltage activated calcium channels [8] affected the gating properties of these membrane proteins.

Voltage activated calcium channels can be classified into two different subfamilies according to their voltage dependence: high voltage- (HVA; named L, N, P/Q, and R-types) and low voltage-activated channels (LVA; named T-type). Regarding the influence of the pore residues on the gating properties of LVA Ca^2+^ channels, a previous report evaluated the role of the selectivity filter residues (EEDD) on the gating properties of T-type Ca^2+^ channels (Ca_V_3.1) by substitution of aspartate by glutamate in the P-loop of domains III (EEED) or IV (EEDE), thus generating a mutant pore closely related to L-type Ca^2+^ channels. In both cases, the L type-like mutant pore affected several gating properties of the Ca^2+^ channel, such as a positive shift of the steady-state activation curve, a change toward a HVA channel [9]. In addition, the value of τ_act_ was increased (slower activation kinetics), whereas the inactivation time constant (τ_inact_) and the deactivation time constant (τ_deact_) values were decreased [10]. The fact that both pore mutations affected the gating properties of the Ca_V_3.1 channels with a similar tendency suggests these two pore locus residues could have an equivalent influence on Ca_V_3.1 channel gating properties.

Recently, we have studied the permeability and selectivity of pore mutant Ca_V_3.1 channels containing glutamate-to-aspartate substitutions in the pore loops of domains I and/or II (named CatSper-like pore mutants: DEDD, EDDD and DDDD) [11]. These mutant Ca_V_3.1 channels presented a reduced sensitivity to inhibition by Cd^2+^, a higher permeability to Cd^2+^, Mn^2+^ and monovalent cations as Na^+^ compare to WT Ca_V_3.1 channels. In the present study, we evaluated gating properties of our pore mutant Ca_V_3.1 Ca^2+^ channels. Our results indicate that substitution of glutamate residues in the pore loops of domains I and II by aspartate residues differentially modify the gating properties of Ca_V_3.1 channel, which implies an asymmetric structure of channel pore in this region.

## 2. Materials and Methods

### 2.1 Cell culture and transfection

Human embryonic kidney (HEK) 293 cells (ATCC, Manassas, VA) were incubated at 37°C in a 5% CO^2^-95% air humidified incubator, in a 35 mm Petri dish contained Advanced Dulbecco’s Modified Eagle Medium (DMEM, Gibco Life Technologies) supplemented with 5% fetal bovine serum (FBS) and 1% penicillin and streptomycin antibiotics. Gene transfer was performed using Lipofectamine and Plus reagent (Invitrogen, Carlsbad, CA) as described previously [11]. Briefly, HEK-293 cells in a 35-mm Petri dish at 60% of confluence (~800 cells/mm^2^) were incubated for 5 hours in 800 μL of serum-free medium containing 100 μL of the transfection solution. Transfection solution was previously prepared with 1.5 µg of the plasmid encoding the WT Ca_V_3.1 calcium channel or the previously reported pore mutant variants (DEDD, EDDD or DDDD) [11], 300 ng of the plasmid encoding the green fluorescent protein (pEGFP, Clontech, CA, USA) and 12 μL of Lipofectamine/Plus reagent, according with the manufacturer’s instructions. After transfection, serum-free medium was substituted for supplemented DMEM and transfected cells were grown at 37 °C, as above-mentioned. 48 h post-transfection, the cell culture was divided to a lower cell density (10% confluence or ~130 cells/mm^2^) onto glass coverslips and 2 h later fluorescent cells were used to perform patch clamp recordings.

### 2.2 Electrophysiological recordings

T-type Ca^2+^ currents were recorded with the classical whole-cell patch-clamp technique [12] at room temperature (20-22°C) using an Axopatch 200B amplifier (Molecular Devices, Sunnyvale, CA). Current data acquisition was performed at 20 KHz and filtered at 5 KHz. During electrophysiological recordings, leak current was subtracted using an on-line P/4 protocol. pClamp 10.6 software (Molecular Devices, Sunnyvale, CA) was used for data acquisition and analysis. The bath solution contained (in mM): 10 CaCl_2_, 125 TEA-Cl, 10 HEPES and 10 glucose (pH 7.3 with TEA-OH). Patch pipettes had a resistance from 2 to 5 MΩ and were filled with an internal solution containing (in mM): 110 CsCl, 5 MgCl_2_, 10 EGTA, 10 HEPES, 4 Na-ATP and 0.1 GTP (pH 7.3 with CsOH). All reagents were purchased from Sigma-Aldrich (St. Luis, Mo, USA).

Current-voltage (I-V) relationships were obtained measuring the peak current amplitude evoked by 200 ms lasting depolarization steps in a range from −80 to +40 mV with a 5 mV increment, from a holding potential (V_h_) of −120 mV.

The steady-state activation of the WT Ca_V_3.1 and the pore mutant channels was approximated from a fit of the amplitudes of the tail currents after a 7 ms depolarizing step to −30 mV, from a V_h_ of −120 mV, following repolarization steps in the voltage range from −80 to +20 mV with a 5 mV voltage increment, to the following equation:

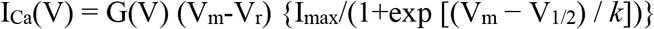

Where I_Ca_ is the measured peak current; G(V) the conductance, which may be voltage dependent; V_m_ is the step potential; V_r_ is the reversal potential; I_max_ is the maximal amplitude of the current; V_1/2_ is the potential of the half-maximal activation and *k* is the slope parameter.

Time constants of activation (τ_act_) and inactivation (τ_inact_) were determined from a fit of current traces with the equation:

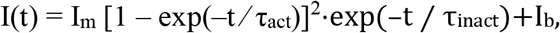

Where I_m_ is a normalization factor and I_b_ is a background component.

The decaying phase of the voltage dependence of τ_act_ (VD τ_act_) was fitted with an exponential function of the form:

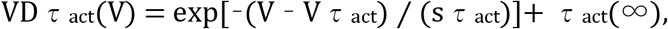

where sτ_act_ is the voltage sensitivity, τ_act_(∞) is the asymptotic value at positive potentials and Vτ_act_ is the voltage at which τ_act_ is equal to 1 + τ_act_(∞).

To assess steady-state inactivation properties we applied a 2-s conditioning pre-pulse to various potentials (from −100 mV to −20 mV), preceded by a test depolarization to −30 mV. Inactivation curves were fitted with the following Boltzmann function:

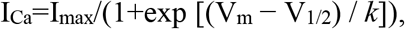

where I_Ca_ is the measured peak current; I_max_ is the maximal amplitude of the current; V_m_ is the test pulse potential; V_1/2_ is the voltage where I_Ca_ has decreased to its half-amplitude and *k* is the slope parameter. In our experimental conditions, the time constant of macroscopic inactivation (τ_inact_) was not voltage dependent in the voltage range from −30 to 0 mV.

Recovery of I_Ca_ from inactivation was studied with a double pulse protocol. After a 200 ms pre-pulse evoked to −30 mV from a V_h_ of −120 mV, we applied a second test pulse (−30 mV) at different time intervals from the pre-pulse. I_Ca_ was normalized by the peak current amplitude obtained with the pre-pulse. In order to estimate the recovery time constant from inactivation (recovery τ), the I_Ca_ fractional recovery was plotted as a function of the interpulse duration and the data were fitted with a single exponential function.

For the deactivation analysis, tail currents were evoked with a 7 ms lasting pre-pulse at −30 mV from a V_h_ of −120 mV; following of 30 ms lasting repolarizing test pulses from −120 mV to −60 mV. Interpulses time lag was at least 800 ms in order to minimize cumulative inactivation. Data were obtained by fitting a curve to the tail-currents following settling of 95% of the membrane capacitance transient from traces. All tail-currents were fitted by a single exponential equation. To characterize the voltage dependence of τ_deact_, we fitted τ_deact_(V) with a growing exponential function in a range of very negative potentials.

### 2.3 Statistical analysis

Electrophysiological data were analyzed and fitted using the Clampfit 10.6 software (Molecular Devices, Sunnyvale, CA). Other curve fitting was done with SigmaPlot 12 software (Systat Software Inc. San Jose, CA, USA). Data were expressed as the mean ± standard error of the mean (SEM). Statistical significance was determined using Student paired *t* test or Analysis of variance (ANOVA) and Tukey’s test for multiple comparisons. A *p*<0.05 was considered significant. Pearson’s correlation analysis was used to compare the activation versus the inactivation kinetics for all Ca_V_3.1 channel versions. For positive correlation coefficients, a Pearson’s correlation coefficient (ρ) >0.95 was considered significant; where *P* values below 0.05 means both parameters tend to increase together. For parameter pairs with negative ρ and *P* values below 0.05, one variable tends to decrease while the other increases. In this case, parameter pairs with *P* values greater than 0.05, mean there is no significant relationship between the two variables. All experiments were repeated at least six times.

### 2.4 Data simulation

Simulations of the kinetic model were made in Java 1.8. We used the library jblas 1.2.4 for all the matrix operations performed for the solution of the differential equations and the Nelder-Mead simplex optimizer implemented in the library Apache Commons Math 3.6.1 for the fittings of the model.

We implemented a real valued error function *E(I)* as a criteria to fit the model with parameters I to the experimental results. We decomposed *E(I)* as the sum of four different functions *E_act_(I)*, *E_inact_(I)*, *E_deact_(I)* and *E_rec_(I)* that represent the mean square error for activation, inactivation, deactivation and recovery from inactivation respectively. The data from the experiments and the simulations was normalized to account for equality between the ranges of the different error functions. Finally, by manual exploration we constructed *E(I)* as the weighted sum of the mean square errors:

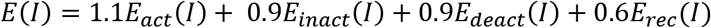

We used the optimizer indicated above to obtain the set of parameters *I* that minimized *E*. Taking into account the convergence of the optimizer into local minima and that this depends on initial conditions, we used the original set of parameters described in [13] and induced random perturbations on all of them, following a normal distribution with mean on the original parameter and with standard deviation similar to 0.2 times its value. With this technique we generated a set of 500 different initial conditions for the optimizer and we selected the set *I* of the optimization that produced the minimum *E(I)*.

### 2.5 Modeling of WT Ca_V_3.1 and pore mutant channels

To better understand our results, we developed a Markovian model with eight states described in the Fig. 6. Although there are many ways to describe the kinetics of T-type channels through mathematical modeling [13–15], in this work we used the model presented in Burgess et al. 2002 [13] because it takes into account many properties of these channels in physiological conditions, such as the gating charges present in transitions between states, as well as their voltage dependency for each subunit [10,13].

We used this model and determined how its parameters had to be modified to fit our own experimental data. Similar to the original model, all of the horizontal transition rates in favor of activation (left to right) and in favor of deactivation (right to left) have a voltage dependence determined by an exponential function. Thus, in the activation direction, the transition rates take the form:

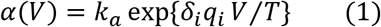

where *k*_*a*_={*k_C1C2_*,*k_C2C3_*,*k_C3O_*,*k_I1I2_*,*k_I2I3_*,*k_I3Io_*}. The parameter *q*_*i*_ (*i*=1,2,3) is the gating charge associated with the *i^th^* box in the model, on each transition *δ_i_* represents the fraction of its voltage dependence towards activation in the *i^th^* box, *V* is the membrane potential fixed by voltage clamp and *T*=25.4 mV represents the thermal energy of room temperature in electron volts.

Similarly, we have the following form for transition rates in the deactivation direction:

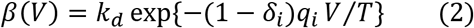

where *k_d_*={*k_C2C1_*,*k_C3C2_*,*k_OC3_*,*k_I2I1_*,*k_I3I2_*,*k_IoI3_*}. The remaining parameters in (2) are the same as in (1).

The model consists of 24 parameters summarized in Table 2 for the WT Ca_V_3.1 and the pore mutant channels. In it *k_C3O_*=*k_I3Io_* and *k_OC3_*=*k_IoI3_*.

The kinetic scheme presented in Fig. 6 can be solved in time by means of the transition matrix *A(V)* which is voltage dependent and is a function of the rate constants *k_a_*,*k_d_*,*δ_i_*,*q_i_*. Let *X*=(*C1*,*C2*,*C3*,*O*,*I1*,*I2*,*I3*,*Io*) be the vector with the values of the eight states, the kinetics of the system over time is described by

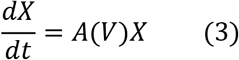

Notice that if *V* is constant, *A(V)* becomes the constant matrix *A_V_* and (3) has an analytical solution so the state vector at time *t* is given by:

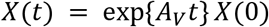

Where *X(0)* represents the state of the system at time 0. In our experiments, the change in membrane potential commanded by the voltage clamp described in our protocols happens so fast that it can be taken as an instantaneous change producing time subintervals where the voltage is constant. In this situation, we solved the equations analytically for each sub interval where the final condition in a subinterval is the initial condition of the next.

Taking into account the previous considerations, we first adjusted the original set of parameters to fit the results obtained for steady state activation, deactivation, inactivation and recovery from inactivation for the EEDD channel. Next, we readjusted the parameters to fit the same results for the mutants DEDD, EDDD and DDDD. The difference obtained in the parameters of each channel can give us insights about their functional properties.

## 3. Results

### 3.1 Differential influence of pore locus residues of domains I and II on the Ca_V_3.1 channel activation kinetics

In the previous work, we prepared three pore mutants DEDD, EDDD and DDDD of Ca_V_3.1 channels and studied primarily their ion conductance properties [11]. The objective of that work was to evaluate the gating properties of those pore mutants. Thus, we first analyzed the voltage dependence of channel activation of the three mutants together with the WT channel using our previously reported Ca^2+^ channels. The three pore mutant channels presented activation curves similar to that of WT Ca_V_3.1 channel (Fig. 1), where the activation threshold voltage was ~−50 mV in all cases. Only the single mutant channel in the domain II (EDDD) presented a half activation voltage (V_1/2_ activation) that was significantly more negative than that of the WT Ca_V_3.1 channel (Fig. 1A, squares); even when the other mutant channels (DEDD and DDDD) also showed the same tendency (Table 1). In addition, the single mutant channel (EDDD) presented a lowest value for the Boltzmann’s constant (*k*=3.3 ± 0.3, Fig. 1B dark gray bar), implying that it has higher voltage dependence than the others. Once again, the other mutant channels (DEDD and DDDD) followed the same tendency but their mean Boltzmann’s constants were not statistically different from that of the WT Ca_V_3.1 channel (Fig. 1B, closed bar).

**Figure 1.**
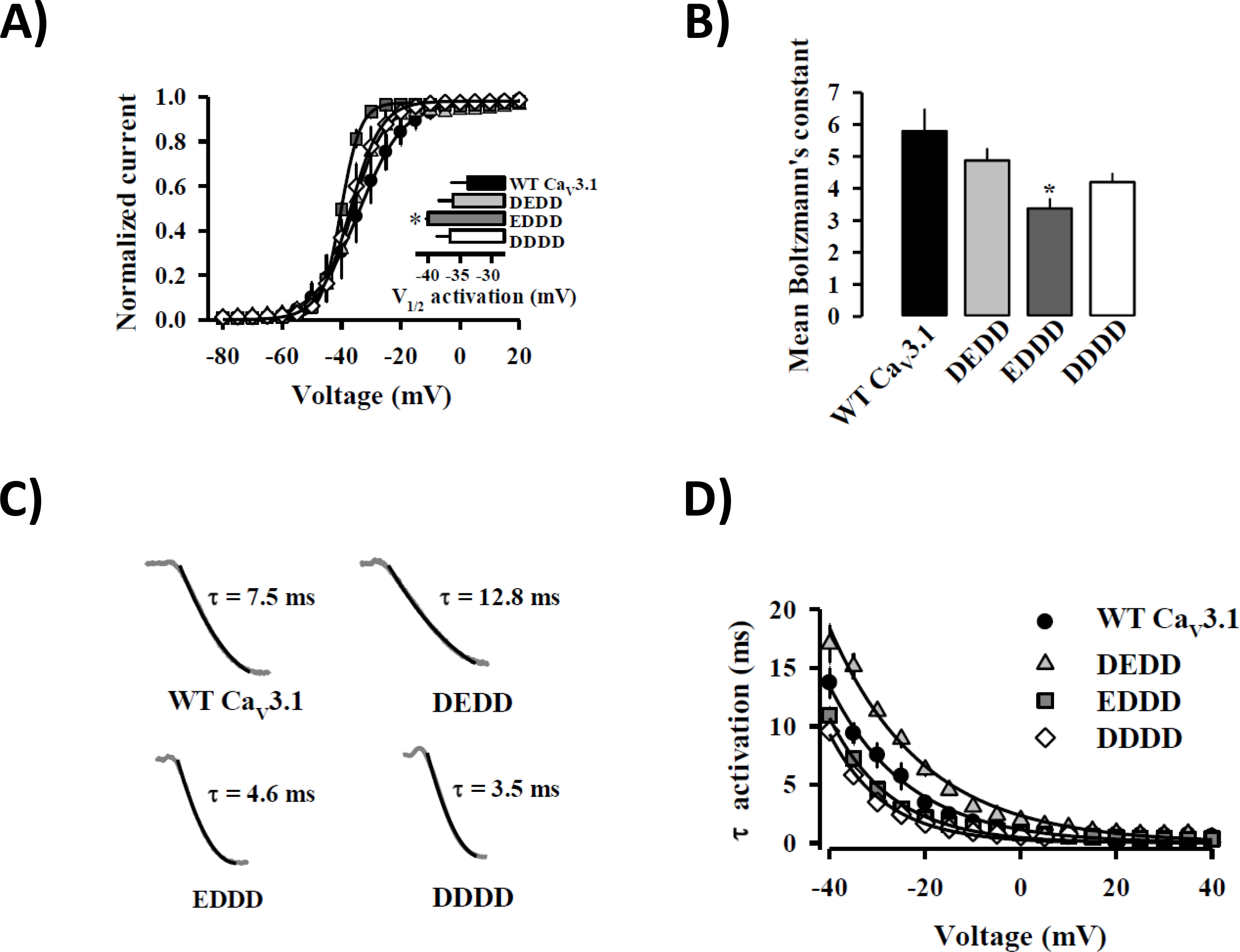
Pore glutamate residue of domain II is important to determine the activation curve parameters and the activation kinetics of the Ca_V_3.1 channel. **A)** Activation curves of WT and mutant Ca_V_3.1 channels. Smooth curves were obtained by fitting data to the Boltzmann function as described in Materials and methods. Only the single mutant Ca_V_3.1 channel (EDDD; squares) presented a statistically left-shifted activation curve from WT channel. Inset: voltage of the half-maximal activation (V_1/2_ act) for all Ca_V_3.1 channel versions; only the V_1/2_ act value for the mutant Ca_V_3.1 channel (EDDD; dark gray bar) was statistically different from WT channel. ***B*)** Consistent with the previously shown activation data, only the mutant Ca_V_3.1 channel (EDDD; dark gray bar) presented a higher voltage sensitivity for activation with a mean Boltzmann’s constant value of 3.3 e-fold/mV. **C)** Representative current traces evoked with a −30 mV test pulse from a V_h_ of −120 mV for the WT Ca_V_3.1 channel and the pore mutant channels (DEDD), (EDDD) and (DDDD). Shown values correspond to the mean τ_act_ for each Ca_V_3.1 channel version. **D)** Voltage dependency of the time constant of activation (τ_act_) for all the Ca_V_3.1 channel versions. The single mutant Ca_V_3.1 channel (DEDD; triangles) showed bigger τ_act_ values than the WT channel (EEDD; circles); whereas both the single mutant Ca_V_3.1 channel (EDDD; squares) and the double mutant Ca_V_3.1 channel (DDDD; diamonds) presented faster activation kinetics than the WT channel. In all cases, symbols and bars represent the mean ± S.E.M. (*n= 6-18 cells*) \**p*< 0.05.

As expected for a higher voltage sensitivity, EDDD mutant channel showed faster activation kinetics than the WT channel, with an activation τ value (τ_act_) of 4.6 ± 0.2 ms at −30 mV (Fig. 1C and D, squares). However, the other mutants displayed a mixed behavior regarding their activation kinetics. The double mutant channel (DDDD) showed the fastest activation kinetics with a τ_act_ value of 3.5 ± 0.1 ms at −30 mV (Fig. 1C and D, diamonds). In contrast, the DEDD mutant channel displayed the slowest activation kinetics (τ_act_ =12.8 ± 0.3 ms at −30 mV, Fig. 1C and D, triangles).

### 3.2 The Ca_V_3.1 channel inactivation kinetics is accelerated by substitution of the pore glutamate residue of domain I

In order to explore whether pore mutations can modify the inactivation process of Ca_V_3.1 channels, we analyzed this parameter. The steady-state inactivation curves showed that only the DEDD mutant channel inactivates at slightly more hyperpolarized voltages (Fig. 2A, triangles); with a half inactivation voltage (V_1/2_ inactivation) of −63 ± 0.6 mV (Figure 2B, light gray bar) compared to that of the WT channel, −61 ± 0.6 mV (Table 1). Despite this discreet effect on the steady-state inactivation curves, the inactivation kinetics was more sensitive to the substitution of the glutamate residues of domain I.

**Figure 2.**
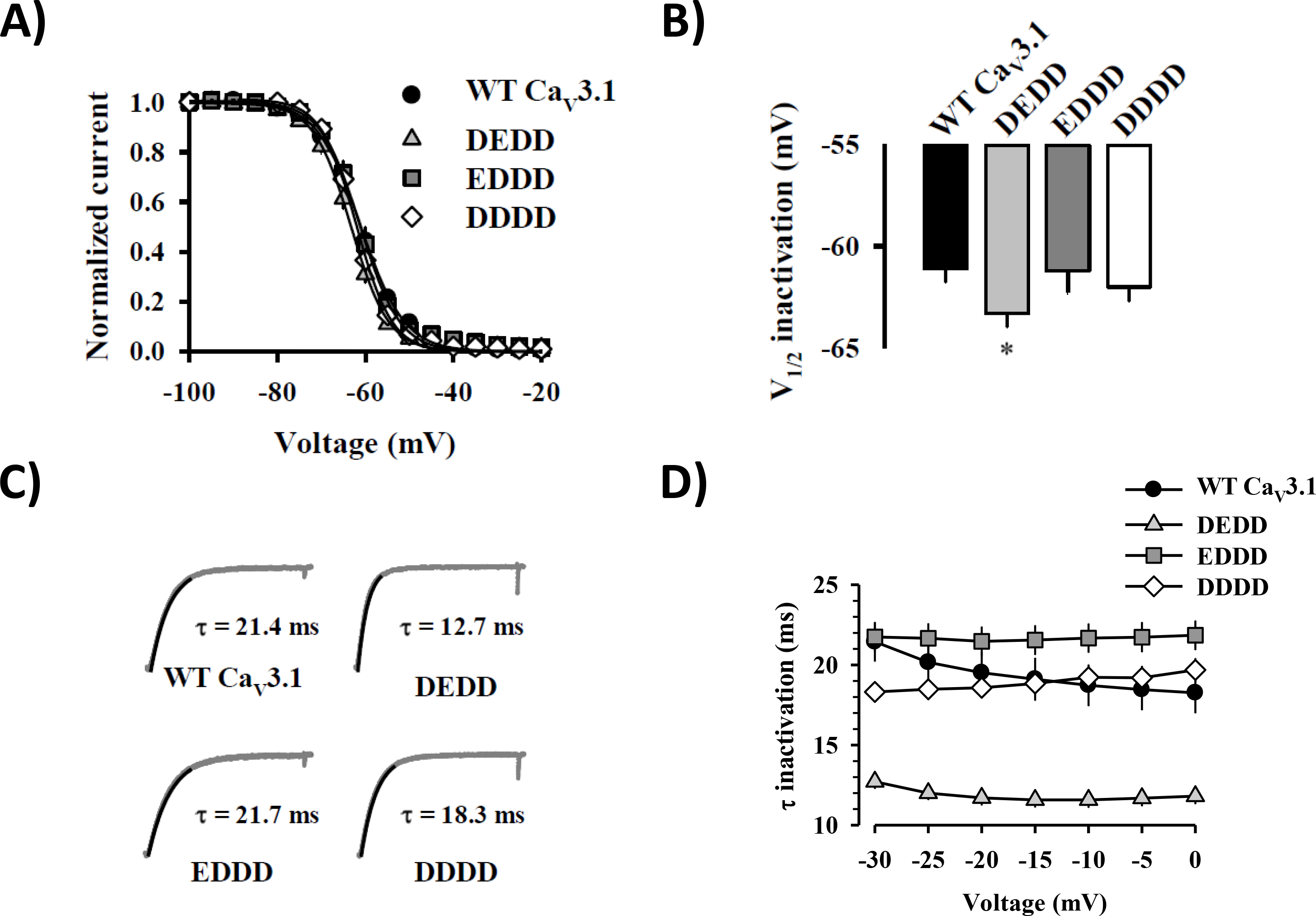
Substitution of pore glutamate residue of domain I slightly left-shifted the steady-state inactivation curve and accelerated the inactivation kinetics of the single mutant Ca_V_3.1 channel (DEDD) ***A*)** Steady-state inactivation curves from all versions of Ca_V_3.1 channel studied. The single mutant Ca_V_3.1 channel (DEDD) was the only channel version with an inactivation curve statistically different from the WT Ca_V_3.1 channel (EEDD). ***B*)** Mean half-voltage of inactivation (V_1/2_ inact) of WT and mutant Ca_V_3.1 channels. Only the V_1/2_ inact of the single mutant Ca_V_3.1 channel (DEDD) was statistically different from WT Ca_V_3.1 channel (−63.3 ± 0.7 versus −61 ± 0.6 mV; respectively). ***C*)** Representative current traces evoked with a −30 mV test pulse from a Vh of −120 mV for the WT Ca_V_3.1 channel and the pore mutant channels (DEDD), (EDDD) and (DDDD). Shown values correspond to the mean τinact for each Ca_V_3.1 channel version. ***D*)** The inactivation kinetics was clearly accelerated in the case of the single mutant Ca_V_3.1 channel (DEDD) compared to WT Ca_V_3.1 channel. In all panels, symbols and bars represent the mean ± S.E.M. (*n=7-11*) \**p*<0.05.

In all Ca_V_3.1 channel versions assayed in this report, the inactivation kinetics were voltage independent in a voltage range from −30 up to 0 mV. However, the inactivation τ value (τ_inact_) was constant and different among the mutant channels. DEDD Ca_V_3.1 channel mutant showed a faster inactivation process, presented τinact constant mean value of 12 ± 0.2 ms (Fig. 2C and D, triangles); whereas DDDD mutant channel showed an statistically similar τ_inact_ constant mean value to WT Ca_V_3.1 channel (19 ± 0.2 vs 20 ± 0.2 ms; Fig. 2C and D, diamonds vs circles respectively). In contrast, substitution of the glutamate residue by aspartate on the pore locus of domain II of Ca_V_3.1 channel (EDDD) slightly slowed the inactivation kinetics, and showed a τ_inact_ mean value of 21.7 ms but it was statistically similar to the τ_inact_ mean value for WT Ca_V_3.1 channel at −30 mV (20 ± 0.2 ms; Fig. 2C and D squares vs circles, respectively).

### 3.3 The activation-inactivation coupling

Previous reports have suggested that the activation and inactivation kinetics are coupled processes in LVA Ca^2+^ channels [15,16]. As a consequence of this phenomenon, the observed change in macroscopic inactivation kinetics induced by point mutations in an ion channel could be a consequence of a modification of the activation kinetics in the first place [16]. In order to evaluate the activation-inactivation relationship in our constructs, we performed a correlation analysis of both parameters according to [17]. As expected, WT Ca_V_3.1 channels (EEDD; circles) showed a linear correlation between its activation and inactivation kinetics (Fig. 3. Correlation parameters *P*<0.001, ρ=0.96), suggesting a good coupling between both processes. Differently, mutant Ca_V_3.1 channels DEDD and EDDD (Fig. 3 triangles and squares, respectively) showed moderated correlations between both parameters (DEDD: *P*<0.01, ρ=0.8; EDDD: *P*=0.001, ρ=0.89). In fact, the mutant Ca_V_3.1 channel (DEDD; Fig. 3 dashed line) displayed a non-linear correlation for its activation and inactivation kinetics, emphasizing a less efficient coupling between both processes. However, DDDD mutant Ca_V_3.1 channel (Fig.3, diamonds) presented the strongest phenotype where no correlation was observed between both parameters (*P*=0.97, ρ=−0.01), suggesting an additive role of these residues in coupling the activation and inactivation processes.

**Figure 3.**
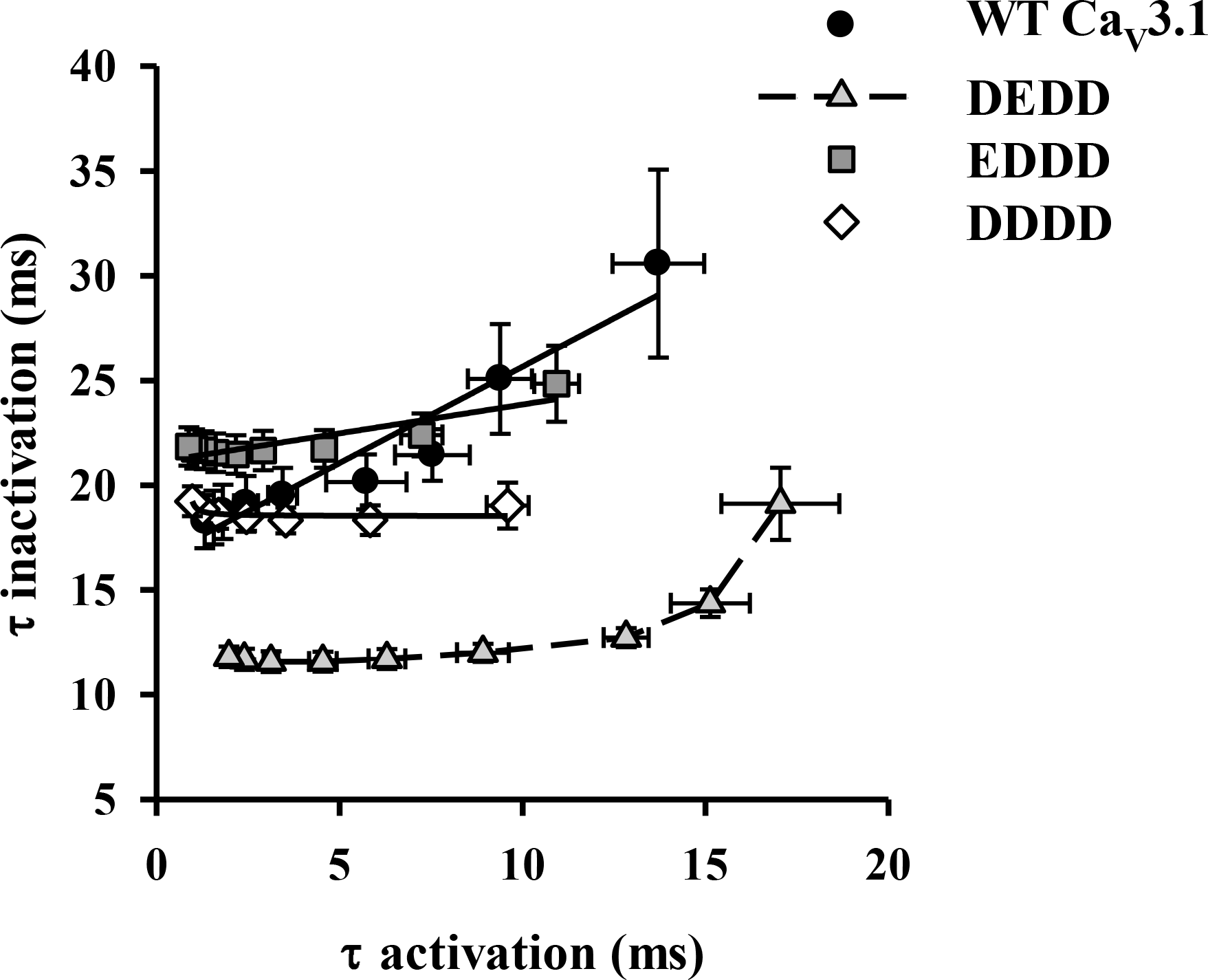
Correlation coefficients *R* and probability values *P* for WT and mutant Ca_V_3.1 channels showed that double pore mutation (DDDD) causes deficient coupling between the activation and inactivation processes. Pearson’s correlation analysis, according to [17], for the activation (τ_act_) and inactivation (τ_inact_) processes of WT (EEDD; circles) and mutant Ca_V_3.1 channels (DEDD, EDDD and DDDD; triangles, squares and diamonds, respectively). Linear correlation parameters for WT *P*<0.001, ρ=0.96; and single mutant Ca_V_3.1 channels (EDDD) *P*=0.001, ρ=0.89. Only the mutant Ca_V_3.1 channel (DEDD; dashed line) showed a non-linear correlation for its activation and inactivation kinetics emphasizing the uncoupling between both processes, where *P*<0.01, ρ=0.8. Double mutant Ca_V_3.1 channel (DDDD) presented the strongest uncoupling between its activation and inactivation kinetics, where *P*=0.97, ρ=-0.01. In all cases, symbols represent the mean ± S.E.M. (*n=6-11*).

### 3.4 The recovery from inactivation was slowed by substitution of the pore glutamate residues of Ca_V_3.1 channel domains I and II

Consistently with a general fast disposition for inactivation for the Ca_V_3.1 mutant channels, their recovery from inactivation was slower as compared to the WT Ca_V_3.1 channel (Fig. 4). Both EDDD and DDDD mutant channels showed a similar recovery time constant (recovery τ), while DEDD mutant channel presented the biggest τ value (590 ms) for the recovery from inactivation (Fig. 4B, triangles and light gray bar in inset).

**Figure 4.**
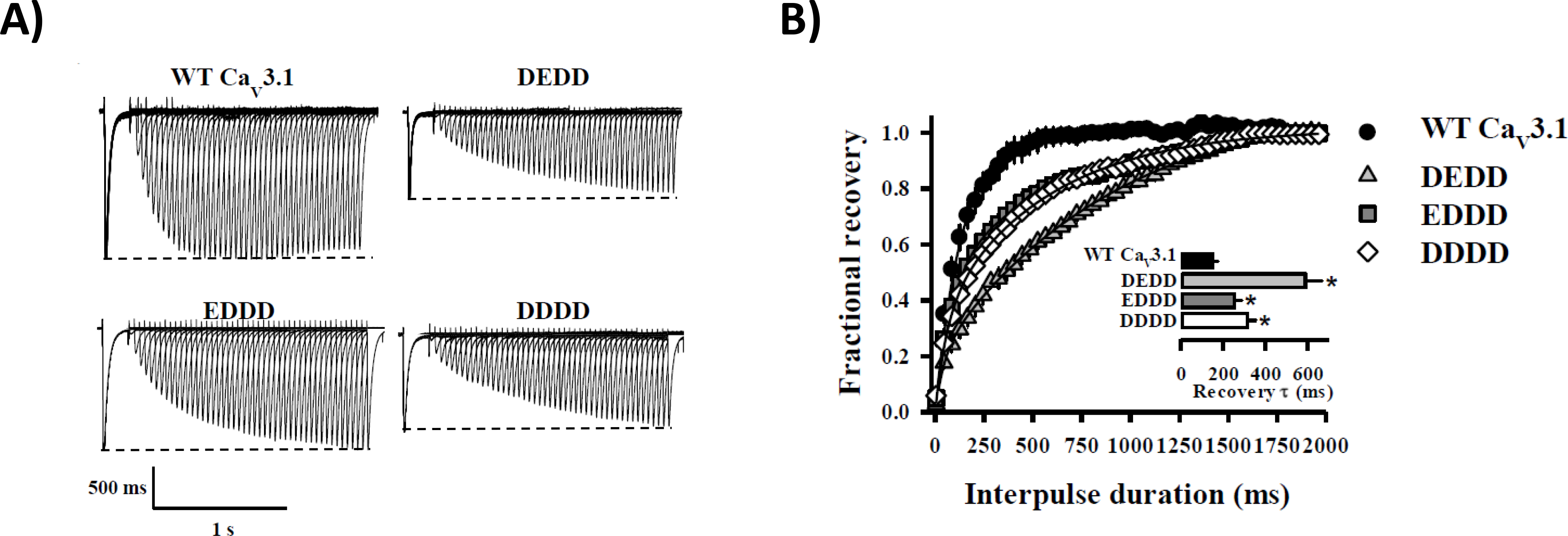
Recovery from inactivation was slowed down in all the mutant Ca_V_3.1 channels. **A)** Representative current traces recorded during the application of the reactivation protocol to HEK-293 cells expressing the WT (EEDD) and pore mutant channels (DEDD, EDDD and DDDD). In all cases, dashed lines represent the initial Ca^2+^ current amplitude. **B)** Reactivation kinetics of the WT and the mutant Ca_V_3.1 channels. Average reactivation curves for WT (EEDD; circles) and mutant Ca_V_3.1 channels (DEDD, EDDD and DDDD; triangles, squares and diamonds, respectively). Continuous lines are the fit of the experimental points as described in Materials and methods. In all cases, mutant Ca_V_3.1 channels presented a slower reactivation kinetics than the WT channel. Inset: mean half-maximal reactivation (recovery τ) for all Ca_V_3.1 channel versions; recovery τ values for all the mutant Ca_V_3.1 channels were statistically different from the WT channel. In all cases, symbols and bars represent the mean ± S.E.M. (*n=8-12*) \**p*<0.05.

### 3.5 The Ca_V_3.1 channel deactivation kinetics accelerates in the double mutant channel

As the last biophysical parameter of the Ca_V_3.1 channels studied here, we evaluated their deactivation kinetics. Only DDDD mutant channel showed a deactivation kinetics statistically faster than the WT Ca_V_3.1 channel (Fig. 5, open diamonds). Despite the DEDD mutant channel presented the same tendency, its deactivation kinetics was not different from that of the WT (Fig. 5, triangles).

**Figure 5.**
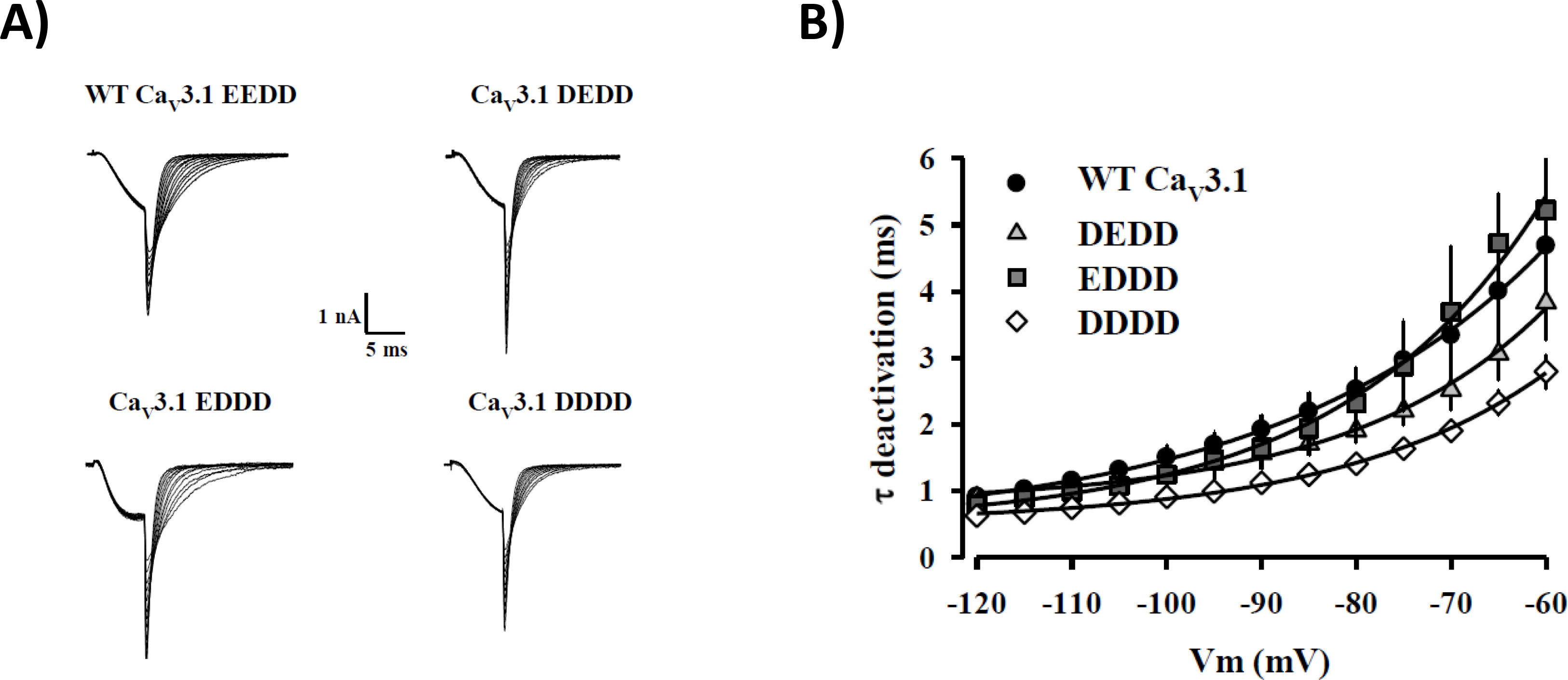
Single (DEDD) and double (DDDD) pore mutant Ca_V_3.1 channels in the selectivity filter showed faster deactivation kinetics than WT channels. **A)** Representative tail current traces recorded with the deactivation protocol to HEK-293 cells expressing the WT (EEDD) and pore mutant channels (DEDD, EDDD and DDDD). **B)** Deactivation curves for WT (EEDD; circles) and mutant Ca_V_3.1 channels (DEDD, EDDD and DDDD; triangles, squares and diamonds, respectively). Continuous lines are the fit of the experimental points. In all cases, symbols represent the mean ± S.E.M. (*n=6-7*) \**p*<0.05.

**Fig. 6.**
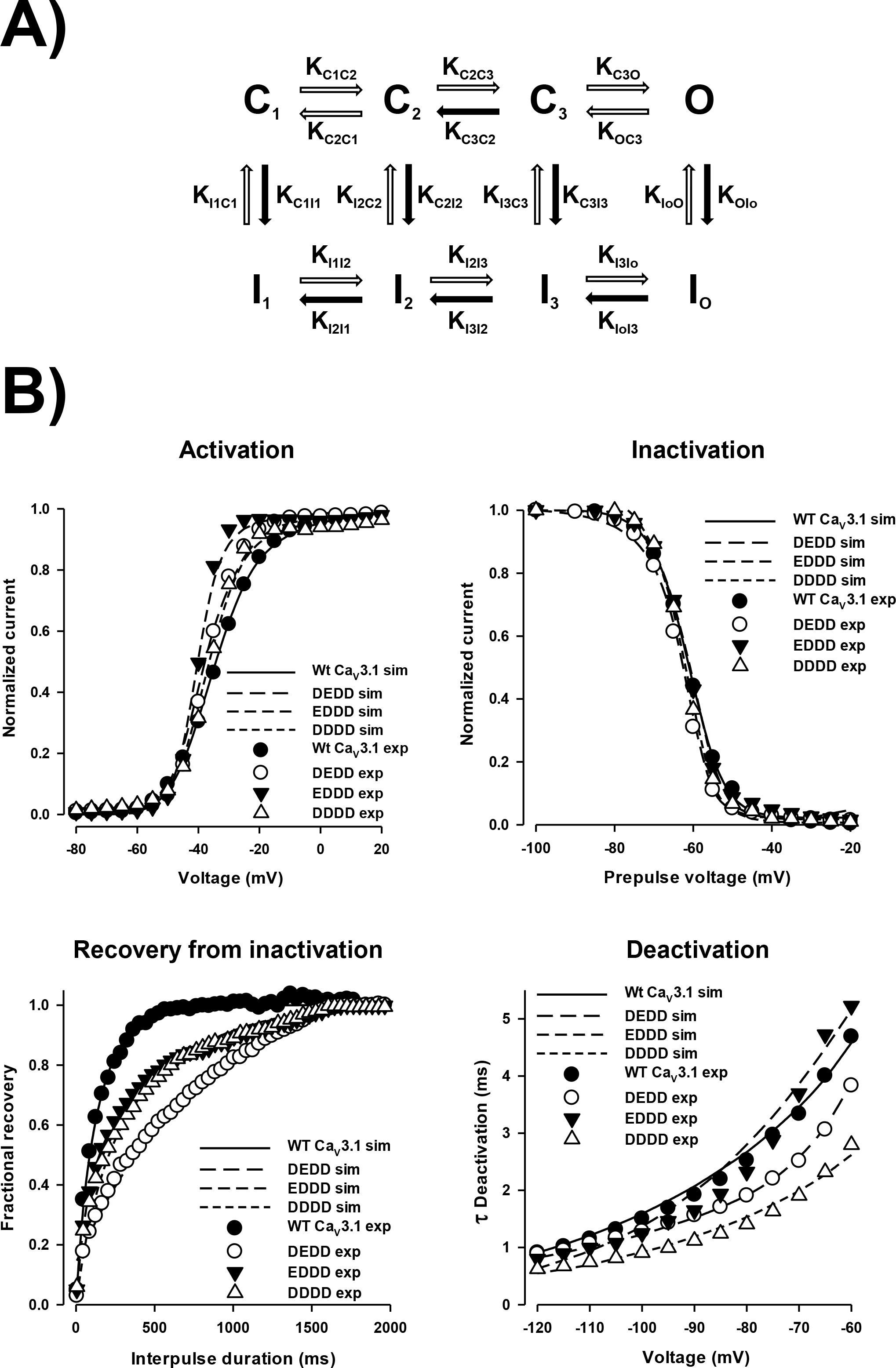
Kinetic model of gating parameters of WT and pore mutant Ca_V_3.1 channels. **A)** At a very negative potential (V_h_= −120 mV), WT Ca_V_3.1 channels are in the resting state, in equilibrium between the closed state C1 and the inactivated state I_1_. After a depolarizing stimulus, Ca_V_3.1 channels undergo a sequence of transitions among closed states (C_1_–C_3_) and eventually transit toward an open state (O), followed by their spontaneous transition to an inactivated state (I_O_). For recovering from inactivation, WT Ca_V_3.1 channels transit among inactivated states and escape from inactivation via closed states, mainly through the I_1_-C_1_ transition. In general, selectivity filter mutations (DEDD, EDDD and DDDD) within the channel pore alter the equilibrium between the inactivated state I_1_ and the closed state C1 of Ca_V_3.1 channels. The decrease in the constant rate of the transition I_1_C_1_ predicts the deficient transit of the channel by its main recovering from inactivation pathway (C_1_-I_1_). Modifications of the transitions among inactivated states (I_1_-I_O_) also contribute to slow down the recovery from inactivation of mutant Ca_V_3.1 channels. Closed arrows represent the voltage-dependent transitions. Open arrows indicate the transitions for which constant rate was significantly increased or decreased by pore mutations within domains I and/or II. **B)** Simulations of biophysical parameters obtained using the kinetic model shown in panel A. Lines superimposed (modeled results) over experimental data (symbols) show accurate fitting among data groups for activation, inactivation, recovery from inactivation and deactivation parameters.

### 3.6 Analyzing domain I and II Ca_V_3.1 pore mutations with an 8-state Markovian scheme

To better understand the impact of pore mutations on Ca_V_3.1 channel kinetics, we used an eight-state Markovian scheme according to [13]. The scheme shown in Fig. 6A could accurately describe many features of the experimental data, including steady-state activation and inactivation curves obtained over a voltage range from −100 to −20 mV (Fig. 6B, upper panels), and recovery from inactivation and deactivation time courses (Fig. 6B, lower panels). Simulated results (lines) were compared with experimental data (symbols) for activation, inactivation, recovery from inactivation and deactivation of WT Ca_V_3.1 and pore mutant channels. For all protocols tested, the data show that the model simulations closely resemble the time course and voltage dependence of WT Ca_V_3.1 and pore mutant currents. The model parameters (Table 2) closely fit the experimental data for the above mentioned biophysical parameters.

In general, pore mutant channels (DEDD, EDDD and DDDD) showed an increase of closed to open channel state transition constant values (K_C1C2_, K_C2C3_, K_C3O_), which is consistent with a slight left shift of pore mutant channel activation curves. On the contrary, transition constant values from close/open to inactive states (K_C3I3_, K_C2I2_, K_C1I1_ y K_OIo_) presented minimal changes compared to control channels, supporting statistically similar steady-state inactivated curves for WT and pore mutant channels. Regarding recovery from inactivation, pore mutant channels (DEDD, EDDD y DDDD) showed a reduction of transition constant values from inactivated to close states (K_I1C1_, K_I2C2_, K_I3C3_). It is worth noting that pore mutations deeply affect the main recovery from inactivation pathway of the Ca_V_3.1 channel, the I1-C1 transition route. These data are consistent with the recorded slower kinetics for recovery from inactivation of the pore mutant channels examined here. At last, transition constant values from open to C3 closed state (K_OC3_) increased for pore mutants DEDD and DDDD, but decreased for pore mutant EDDD. These data partially explain faster deactivation of pore mutants DEDD and DDDD (Table 2).

## 4. Discussion

Previous reports have evaluated the role of the selectivity filter residues (EEDD) on the gating properties of T-type Ca^2+^ channels substituting aspartate by glutamate in the P-loop of domains III (EEED) or IV (EEDE), thus generating mutant pore loops which are closely related to L-type Ca^2+^ channels. Both L type-like pore mutations affected several gating properties with respect to wild-type Ca_V_3.1 channels and produced a series of similar effects (shown in Table 1). These effects include: a positive shift of the steady-state activation curve [9], an increase in τ_act_ value (slower activation kinetics), a decrease in the τ_inact_ and τ_deact_ values (faster kinetics), without affecting the reactivation or recovery from inactivation of the Ca^2+^ current [10]. The fact that both pore mutations, in the P-loop of domains III (EEED) or IV (EEDE), affected the gating properties of the Ca_V_3.1 channels with a similar tendency suggests pore locus residues could have an equivalent influence on Ca_V_3.1 channel gating properties. However, the role of pore residues in the P-loop of domains I or II on the Ca_V_3.1 channel gating properties had not been assessed.

In the present report we studied the gating properties of Ca_V_3.1 channel mutants in the P-loop of domains I (DEDD), II (EDDD) or both (I and II, DDDD), generating mutant pore loops closely related to CatSper-type Ca^2+^ channels, which were originally prepared to study ion conductance [11]. CatSper is a Ca^2+^ channel specific of the male germ cell line which is involved in sperm hyperactivation, a vigorous and asymmetric flagellar beat which is necessary for sperm to reach the homologous oocyte within the female genital tract [18]. The CatSper selectivity filter has a pore sequence constituted by four aspartate residues (DDDD) which is highly Ca^2+^ selective. It is worth noting that it has not been possible to heterologously express CatSper. However, this ion pore is more closely related to the K^+^ channel pore than to the Ca^2+^ channel pore because it is structurally formed by four independently-encoded α1 subunits [19].

### 4.1 The four pore residues of Ca_V_3.1 channel differentially contribute to its gating properties

As expected, CatSper-like pores affected several gating properties of the Ca_V_3.1 channel, but the observed effects on its biophysical parameters depend on the mutated domain (shown in Table 1). In general, single (DEDD, EDDD) or double (DDDD) mutants presented different activation time constant (τ_act_) values. Whereas the DEDD Ca_V_3.1 mutant channel showed slower activation kinetics than the WT Ca_V_3.1 channel, the EDDD and the DDDD mutant channels presented a faster activation process (Fig. 1C). According to our results, the glutamate pore residue in domain I could have a preponderant role in the activation kinetics of Ca_V_3.1 channels since a point mutation of this residue is enough to slow down this process. A previous report showed that the substitution of aspartate residues by glutamate in the pore locus of domains III or IV accelerate the Ca_V_3.1 activation kinetics [10], which points out to the particular influence of the glutamate residue in the pore locus of domain I in the Ca_V_3.1 activation kinetics (Table 1). In addition, the substitution of the glutamate pore residue of domain I altered in some way the activation-inactivation kinetic coupling of the Ca_V_3.1 channel. As previously reported [15], the macroscopic inactivation of Ca_V_3.1 Ca^2+^ channels is allosterically coupled to the movement of at least the three first-activated voltage sensors. Simplifying this concept, it means that to a fast activation kinetics corresponds a fast macroscopic inactivation. In our hands, DEDD mutant channel showed slow activation kinetics but the fastest macroscopic inactivation (Fig. 1D vs 2D, triangles). This phenotype could be explained by a stabilization of the inactive and/or closed states of Ca_V_3.1 channel, which could retard the channel transition from the closed to the open state, possibly increasing the latency to the first opening of the single channel. According to our modeled results (Table 2), all pore mutant channels (DEDD, EDDD y DDDD) showed a reduction of transition constant values from inactivated to close states (K_I1C1_, K_I2C2_, K_I3C3_). Therefore, the most plausible hypothesis to explain the slow activation kinetics phenotype of DEDD mutant channel is stabilization of its inactive state more than a deficient transition among its closed states. However, single channel analysis is necessary to confirm our results presented here.

In addition, the modifications in the gating properties of DEDD mutant Ca_V_3.1 channel here reported, allowed us to explain in part the strong reduction in the Ca^2+^ current density shown previously for this mutant channel [11]. Primarily, the combination of a slow activation kinetics (Fig. 1, triangles) with a fast inactivation kinetics (Fig. 2, triangles) should reduce the peak macroscopic Ca^2+^ current. Furthermore, the slow recovery from inactivation could leave a significant fraction of inactivated channels during the current measurements in our previous report.

Regarding EDDD mutant channel, substitution of the glutamate pore residue of domain II by an aspartate seems to destabilize the Ca_V_3.1 channel’s close state, accelerating its transition to the open state (Fig. 1, squares), slowing its inactivation kinetics and having mild effects on the channel’s deactivation kinetics (Table 1). These results are consistent with previous observations where a left shift of the activation curve is associated with slow deactivation kinetics in a LVA mutant L-type Ca^2+^ channel [20]. Interestingly, a recent report pointed out the relevance of the glutamate pore residue of domain II of CaV1.2 on Ca^2+^-dependent inactivation (CDI) of HVA Ca^2+^ channels [21], where the selectivity filter forms a CDI regulatory gate, suggesting a selectivity filter contribution to the inactivation paradigm of HVA Ca^2+^ channels. Our results indicate that the glutamate pore residue of domain II also contributes regulating voltage-dependent inactivation of LVA Ca^2+^ channels.

### 4.2 The pore filter residues of Ca_V_3.1 channel and the S5 and S6 transmembrane segment movement, a speculative hypothesis

Taking all our results into account, we can conclude that every residue within the pore loop of the Ca_V_3.1 channel differentially influences its gating properties. This differential influence of the pore residues on channel gating properties could be related in principle with the Ca_V_3.1 pore asymmetry. Unfortunately, the LVA Ca^2+^ channel crystal structure still remains elusive to support our results with structural data, but considering the recent structural information obtained for the Ca_V_1.1 Ca^2+^ channel complex [22], this hypothesis is highly probable. In this elegant study, Wu and collaborators showed that the Ca_V_1.1 pore domain presents a pseudo-symmetry, where the external loop which binds the S5 transmembrane segment with the P1 α helix and the external loop which binds the P2 α helix and the S6 transmembrane segment of each channel repeat present marked differences in primary sequences and conformations. Moreover, in the same study Wu et al. (2015) [22] indicated that the S5 and S6 segments of the four repeats exhibit slightly different orientation angles and the S4-S5 linker helices deviate from each other to a different extent, increasing the Ca_V_1.1 channel asymmetry. Considering this structural information, we can speculate that the Ca_V_3.1 pore domain could also present conformational differences among its S5 and S6 transmembrane segments, among its S4-S5 linker helices and the external loops responsible for the binding of P1 and P2 α helices with the S5 and S6 transmembrane segments, respectively. Indeed, LVA Ca^2+^ channels contain a large conserved external loop (residues 211-336 for Ca_V_3.1 channel) between the S5 transmembrane segment and the P pore loop of domain I [17], which is different in length with respect to the other three external loops and supports the pseudo-symmetry in LVA channels. At least two different reports have explored the role of pore-lining S6 segment residues in Ca_V_1.2 and Ca_V_3.2 channels [20,23]. The distal S6 transmembrane segment seems to be especially important for the voltage sensitivity of activation and the activation and deactivation kinetics of Ca_V_1.2 channel; substitution of S6 pore region residues of domain I (IS6 L(434), domain II (IIS6; I781) or domain III (IIIS6; G1193) transformed the Ca_V_1.2 from a HVA into a LVA Ca^2+^ channel, probably for altering backbone-backbone helix interactions in a hydrophobic environment (reviewed in [20]). In the case of Ca_V_3.2 channels, the S4-S5 helix linker of domain II (IIS4-S5) interacts with the distal S6 pore region of domains II (IIS6) and III (IIIS6). In particular residues V907 and T911 of S4-S5 helix linker of domain II (IIS4-S5) interact with S6 of domains II (IIS6; I1013) and III (IIIS6; N1548) demonstrating that S4-S5 and S6 helices from adjacent domains are coupled in LVA Ca^2+^ channels [23].

Regarding the influence of pore loop on the gating properties of LVA Ca^2+^ channels, a previous report demonstrated that cysteines in the external IS5-P connecting loop are involved in the voltage dependency of activation and inactivation kinetics of Ca_V_3.1 channel [17]. In the same report, the authors demonstrated that four out of six cysteines of the IS5-P loop produced no functional Ca^2+^ channels, even when mutants reached the plasma membrane, whereas the altered gating properties of mutant channels in cysteines C289 and C313 demonstrated their participation in activation, inactivation and deactivation of Ca_V_3.1 channels [17].

Our results indicate that in addition to channel ion selectivity and permeability [11], deep pore residues within the selectivity filter affect different biophysical parameters of the Ca_V_3.1 channel (Tables 1 and 2). Considering that LVA and HVA Ca^2+^ channel are structurally close-related [22], it is quite possible that each pore residue in the Ca_V_3.1 channel selectivity filter is also differently orientated. Thus, the substitution of a specific pore residue by a different amino acid probably changes its structural orientation angle and could have differential repercussions on the gating properties of the Ca_V_3.1 channel for allosterically altering the movement of amino acid residues from the P1 and/or P2 α helices in LVA Ca^2+^ channels. Subsequently this change in the movement of P1 and/or P2 α helices could affect the displacement of S5 or S6 transmembrane segments in response to the mechanical stimulus of S4-S5 linker helix after a membrane depolarization. At least for CNG1 channels, it has been suggested that its putative P helix rotates around its long axis during the channel gating [24]. Moreover, the W434F substitution within the pore helix causes permanent C-type inactivation of Shaker K^+^ channel as a consequence of the interaction among aromatic residues in the P helix with amino acids in the selectivity filter, which hold the selectivity filter open, and leads to inactivation of the channel [25]. In the case of the Ca_V_1.2 channel pore domain, P1 and P2 α helices which flank the selectivity filter, seem to be overlaid to the S5 and S6 transmembrane segments, respectively [22]. Taking into account the previous references, potential interactions between amino acid residues of P1 and P2 α helices, the pore selectivity filter and amino acid residues from S5 and S6 transmembrane segments of Ca_V_3.1 channels cannot be discarded. However, a correlation mutation analysis among the selectivity filter residues of Ca_V_3.1 channel and some residues from its P1 and/or P2 α helices is necessary to confirm our above-mentioned hypothesis.

## 5. Conclusions

Pore selectivity filter residues (EEDD) differentially contribute to the Ca_V_3.1 channel gating properties, suggesting a pore pseudo-symmetry in LVA Ca^2+^ channels, as described for HVA Ca^2+^ channels. Briefly, the substitution of the glutamate pore residue by aspartate within domain I (DEDD) affects the coupling between activation and inactivation processes, resulting in a more stable inactivation state and a slower recovery from inactivation without no change in the deactivation kinetics. In contrast in domain II, the single mutant channel (EDDD) presented a less stable close state, allowing an easier channel transition to the open state with inactivation kinetics similar to WT Ca_V_3.1 channels, and a slower deactivation kinetics. Double mutant channels in domains I and II (DDDD) presented less coupled activation and inactivation processes; faster activation, inactivation and deactivation kinetics and slower deactivation process than WT Ca_V_3.1 channel. These phenotypes could be explained by potential interactions between P1 and P2 α helices amino acid residues, pore selectivity filter and/or amino acid residues from S5 and S6 transmembrane segments of Ca_V_3.1 channels.

## 6. Acknowledgements

Authors thank the technical support of Carmen Santana-Calvo in the engineering of the mutant Ca_V_3.1 channels.

## 9. Author contribution

EGL performed and analyzed the electrophysiological experiments. TN conceived and designed the mutagenesis of Ca_V_3.1 channel. AS and AD modeled and simulated the biophysical parameters of Ca_V_3.1 channel kinetics model. ILG conceived and coordinated the entire study. All authors contributed to the paper writing and approved the final version of the manuscript.

---

Table 1 Summary of gating properties of the mutant Ca_V_3.1 channels (DEDD, EDDD and DDDD) compared with the WT channel

Table 2 Biophysical parameters obtained by simulating the experimental data of Ca_V_3.1 wild type channel and the DEDD, EDDD and DDDD pore mutants

## References

[1] W.A. Catterall, N. Zheng, Deciphering voltage-gated Na^+^ and Ca^2+^ channels by studying prokaryotic ancestors, Trends Biochem. Sci. 40 (2015) 526–534. doi:10.1016/j.tibs.2015.07.002.

[2] W.A. Catterall, Ion channel voltage sensors: Structure, function, and pathophysiology, Neuron. 67 (2010) 915–928. doi:10.1016/j.neuron.2010.08.021.

[3] P. Proks, C.E. Capener, P. Jones, F.M. Ashcroft, Mutations within the P-loop of Kir6.2 modulate the intraburst kinetics of the ATP-sensitive potassium channel, J. Gen. Physiol. 118 (2001) 341–353. doi:DOI 10.1085/jgp.118.4.341.

[4] G. Yellen, The voltage-gated potassium channels and their relatives., Nature. 419 (2002) 35–42. doi:10.1038/nature00978.

[5] S. Sheng, J. Li, K.A. McNulty, T. Kieber-Emmons, T.R. Kleyman, Epithelial sodium channel pore region. Structure and role in gating, J. Biol. Chem. 276 (2001) 1326–1334. doi:10.1074/jbc.M008117200.

[6] F.J.P. Kúhn, N.G. Greeff, Mutation D384N alters recovery of the immobilized gating charge in rat brain IIA sodium channels, J. Membr. Biol. 185 (2002) 145–155. doi:10.1007/s00232-001-0122-1.

[7] G.E. Flynn, J.P. Johnson, W.N. Zagotta, Cyclic nucleotide-gated channels: shedding light on the opening of a channel pore., Nat. Rev. Neurosci. 2 (2001) 643–651. doi:10.1038/35090015.

[8] A. Yatani, A. Bahinski, M. Wakamori, S. Tang, Y. Mori, T. Kobayashi, A. Schwartz, Alteration of channel characteristics by exchange of pore-forming regions between two structurally related Ca2+ Channels, Mol. Cell. Biochem. 140 (1994) 93–102. doi:10.1007/BF00926748.

[9] K. Talavera, M. Staes, A. Janssens, N. Klugbauer, G. Droogmans, F. Hofmann, B. Nilius, Aspartate Residues of the Glu-Glu-Asp-Asp (EEDD) Pore Locus Control Selectivity and Permeation of the T-type Ca2+ Channel?? 1G, J. Biol. Chem. 276 (2001) 45628–45635. doi:10.1074/jbc.M103047200.

[10] K. Talavera, A. Janssens, N. Klugbauer, G. Droogmans, B. Nilius, Pore structure influences gating properties of the T-type Ca2+ channel alpha1G., J. Gen. Physiol. 121 (2003) 529–40. doi:10.1085/jgp.200308794.

[11] E. Garza-López, J.C. Chávez, C. Santana-Calvo, I. López-González, T. Nishigaki, Cd2+ sensitivity and permeability of a low voltage-activated Ca2+ channel with CatSper-like selectivity filter, Cell Calcium. (2016). doi:10.1016/j.ceca.2016.03.011.

[12] O.P. Hamill, A. Marty, E. Neher, B. Sakmann, F.J. Sigworth, Improved patch-clamp techniques for high-resolution current recording from cells and cell-free membrane patches, Pflugers Arch. Eur. J. Physiol. Arch. Eur. J. Physiol. 391 (1981) 85–100. doi:10.1007/BF00656997.

[13] D.E. Burgess, O. Crawford, B.P. Delisle, J. Satin, Mechanism of inactivation gating of human T-type (low-voltage activated) calcium channels, Biophys. J. 82 (2002) 1894–1906. doi:10.1016/S0006-3495(02)75539-2.

[14] X.J. Wang, J. Rinzel, M.A. Rogawski, A model of the T-type calcium current and the low-threshold spike in thalamic neurons., J. Neurophysiol. 66 (1991) 839–50. doi:10.1152/jn.1991.66.3.839.

[15] J.R. Serrano, E. Perez-Reyes, S.W. Jones, State-dependent inactivation of the alpha1G T-type calcium channel., J. Gen. Physiol. 114 (1999) 185–201. doi:10.1085/jgp.114.2.185.

[16] K. Talavera, B. Nilius, Biophysics and structure-function relationship of T-type Ca2+ channels, Cell Calcium. 40 (2006) 97–114. doi:10.1016/j.ceca.2006.04.013.

[17] M. Karmazinova, S. Beyl, A. Stary-Weinzinger, C. Suwattanasophon, N. Klugbauer, S. Hering, L. Lacinova, Cysteines in the loop between IS5 and the pore helix of Ca_V_3.1 are essential for channel gating, Pflugers Arch. Eur. J. Physiol. 460 (2010) 1015–1028. doi:10.1007/s00424-010-0874-5.

[18] K. Miki, D.E. Clapham, Rheotaxis guides mammalian sperm, Curr. Biol. 23 (2013) 443–452. doi:10.1016/j.cub.2013.02.007.

[19] B.J. Liebeskind, D.M. Hillis, H.H. Zakon, Independent acquisition of sodium selectivity in bacterial and animal sodium channels, Curr. Biol. 23 (2013) R948–R949. doi:10.1016/j.cub.2013.09.025.

[20] S. Beyl, K. Depil, A. Hohaus, A. Stary-Weinzinger, E. Timin, W. Shabbir, M. Kudrnac, S. Hering, Physicochemical properties of pore residues predict activation gating of CaV1.2: A correlation mutation analysis, Pflugers Arch. Eur. J. Physiol. 461 (2011) 53–63. doi:10.1007/s00424-010-0885-2.

[21] F. Abderemane-Ali, F. Findeisen, N.D. Rossen, D.L. Minor, A Selectivity Filter Gate Controls Voltage-Gated Calcium Channel Calcium-Dependent Inactivation, Neuron. (2019) 1134–1149. doi:10.1016/j.neuron.2019.01.011.

[22] J. Wu, Z. Yan, Z. Li, C. Yan, S. Lu, M. Dong, N. Yan, Structure of the voltage-gated calcium channel Cav1.1 complex, Science (80-.). 350 (2015) aad2395–aad2395. doi:10.1126/science.aad2395.

[23] P.O. Demers-Giroux, B. Bourdin, R. Sauvé, L. Parent, Cooperative activation of the T-type Cav3.2 channel interaction between domains II and III, J. Biol. Chem. 288 (2013) 29281–29293. doi:10.1074/jbc.M113.500975.

[24] J. Liu, S.A. Siegelbaum, Change of pore helix conformational state upon opening of cyclic nucleotide-gated channels, Neuron. 28 (2000) 899–909. doi:10.1016/S0896-6273(00)00162-8.

[25] D.A. Doyle, J.M. Cabral, R.A. Pfuetzner, A. Kuo, J.M. Gulbis, S.L. Cohen, B.T. Chait, R. MacKinnon, The structure of the potassium channel: molecular basis of K+ conduction and selectivity., Science. 280 (1998) 69–77. doi:10.1126/science.280.5360.69.

